# Satelight: Self-Attention-Based Model for Epileptic Spike Detection from Multi-Electrode EEG

**DOI:** 10.1101/2021.06.17.448793

**Authors:** Kosuke Fukumori, Noboru Yoshida, Hidenori Sugano, Madoka Nakajima, Toshihisa Tanaka

## Abstract

**Objective:** Because of the lack of highly skilled experts, automated technologies that support electroencephalogram (EEG)-based in epilepsy diagnosis are advancing. Deep convolutional neural network-based models have been used successfully for detecting epileptic spikes, one of the biomarkers, from EEG. However, a sizeable number of supervised EEG records are required for training.

**Approach:** This study introduces the Satelight model, which uses the self-attention (SA) mechanism. The model was trained using a clinical EEG dataset labeled by five specialists, including 16,008 epileptic spikes and 15,478 artifacts from 50 children. The SA mechanism is expected to reduce the number of parameters and efficiently extract features from a small amount of EEG data. To validate the effectiveness, we compared various spike detection approaches with the clinical EEG data.

**Main results:** The experimental results showed that the proposed method detected epileptic spikes more effectively than other models (accuracy = 0.876 and false positive rate = 0.133).

**Significance:** The proposed model had only one-tenth the number of parameters as the other effective model, despite having such a high detection performance. Further exploration of the hidden parameters revealed that the model automatically attended to the EEG’s characteristic waveform locations of interest.

## 1. Introduction

Epilepsy is a chronic brain condition that results in seizures characterized by disorientation and convulsions associated with excessive electrical stimulation of brain neurons [1]. Epilepsy affects about 50 million people worldwide, and there is a severe shortage of specialists who treat epilepsy (epileptologists). For example, Japan only has 700 epileptologists, regardless of having one million patients suffering from epilepsy [2].

Furthermore, a lot of epileptologists’ work is necessary to care for each patient because the diagnosis of epilepsy requires specialized knowledge or skill. This has inspired the development of an automated diagnostic tool to aid epileptologists.

A crucial diagnostic step is the measurement and analysis of an electroencephalogram (EEG). This is because the epilepsy symptoms are determined based on a crucial biomarker called epileptiform discharges (epileptic spikes), which are often seen in a patient’s interictal EEG [3]. Because locating these epileptic spikes is difficult, numerous automated detection techniques have significantly improved over time to support this process[4, 5].

Particularly, neural network (NN)-based techniques have demonstrated high performance in spike detection tasks [6, 7, 8, 9, 10]. Epileptic spikes have been detected on a single-electrode EEG. Although the shape of the epileptic waveforms is similar among patients, each patient’s distribution of the waveforms is unique. To identify these individual differences in distribution among patients, specialists use several montages in epilepsy diagnosis, like bipolar and monopolar, to observe EEGs. This is because the specialists use the EEG of the surrounding or all electrodes to identify epileptic spikes. Therefore, to effectively use the detection methods from a single-electrode EEG, the appropriate montage must be determined manually, limiting its versatility. These facts have influenced studies on machine learning-based automatic spike detection. Since 2019, numerous related studies have used multi-electrode EEG to detect the location of epileptic spikes[11, 12, 13, 14, 15].

For detecting spikes using these multi-electrode EEG, convolutional NN (CNN)- based models, such as the model by Thomas et al.’s [14] and SpikeNet [12], are some of the effective methods. Thomas et al. [14] used CNN to detect epileptic spikes from multi-electrode EEG by calculating the probability of the appearance of epileptic spikes for individual electrodes and then obtaining the maximum of the output probabilities for all electrodes. SpikeNet [12] directly outputs a predicted value from a multi-electrode EEG segment. This was achieved using a combination of deep convolution layers in the spatial and temporal directions. SpikeNet efficiently extracts features using a top convolutional layer that merges the relationships between electrodes and 22 temporal convolutional layers. According to Thomas et al. [12], SpikeNet outperforms the detection performance of the commercially available software, Persyst13 [16], in spike detection.

However, training CNN-based models with many parameters necessitates a large labeled dataset, and gathering a significant amount of epileptic EEG is challenging [17]. Therefore, developing a lightweight model suitable for EEG analysis can contribute to epileptic spike detection from multi-electrode EEG. Here, the analysis of a multi-dimensional time series can consider spike detection from a multi-electrode EEG. According to Farsani et al. [18], the number of parameters in time-series data analysis can be reduced using a mechanism called self-attention (SA). By learning the relationship between all sampled points in a segment, the SA mechanism can efficiently discover significant features. Many convolution and pooling layers are required to implement this with CNNs. In this study, we hypothesized that the trainable parameters in the model for epileptic spike detection from EEG would be reduced using the SA mechanism. To this end, we constructed a spike detection model for multi-electrode EEG called the *self-attention-based lightweight model* (Satelight), which is built with a minimal set of parameters. For the verification of this method, different spike detection approaches were compared using a dataset consisting of clinical EEGs recorded from 50 children with either childhood epilepsy with centro-temporal spikes (CECTS) [19] or focal epilepsy, with supervised labels indicating the location of epileptic spikes annotated by five experts.

## 2. Method

### 2.1. Dataset

Table 1 summarizes the dataset. Interictal EEG recordings were collected from 50 children (23 males and 27 females) with either CECTS or focal epilepsy [19] at the Department of Pediatrics, Juntendo University Nerima Hospital. The patients’ ages at the time of the examination ranged between three and twelve years. The data were recorded from 16 electrodes with the International 10–20 system using the Nihon Koden EEG-1200 system. The sampling frequency was 500 Hz. This dataset was recorded and analyzed with the approval from the Juntendo University Hospital Ethics Committee and the Tokyo University of Agriculture and Technology Ethics Committee.

**Table 1.**
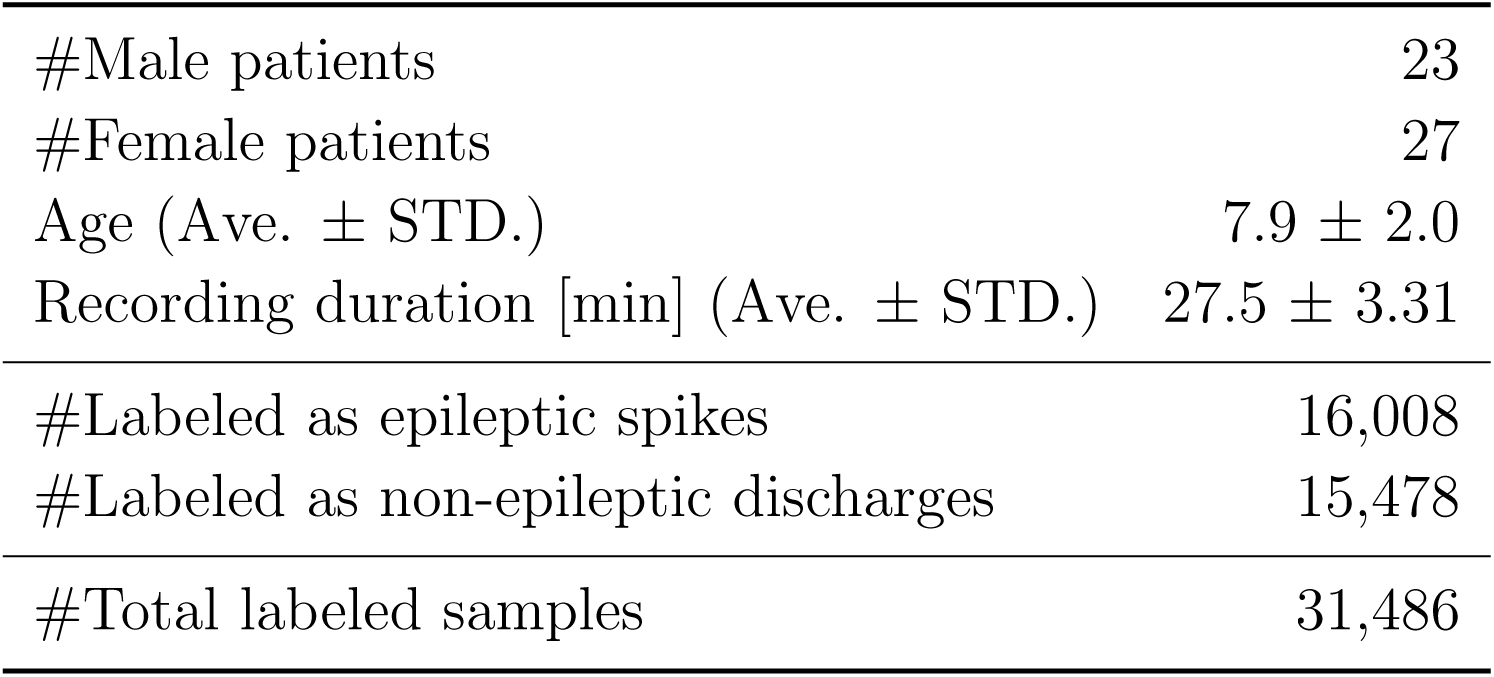
Dataset summary of 50 epileptic EEG. Two neurosurgeons, two clinical technologists, and one pediatrician labeled this dataset. The total number of labeled samples is 31,486.

Measurement EEG recordings were pre-processed as follows. First, using Scipy [20], a peak search method was used to find the EEG’s peak waveforms (minima and maxima)with. With a minimum distance of 100 points, this function extracts both the upward and downward peaks. Meaningless peaks caused by noise and other factors were removed using a threshold set at the 80th percentile of all peaks in one electrode. Second, for the peaks (namely, candidate spikes) that were actual epileptic spikes (spike or spike-and-wave), five experts annotated labels as epileptic. To produce accurate non-epileptic segments, the experts also labeled non-epileptic discharges. These expert-selected non-epileptic discharges, which were mistakenly detected as candidate spikes, included electromyograms, ambient noise, and alpha EEG. Finally, the EEG recordings were cropped every 1.024 s (512 sampling points) from the beginning of the recordings, independent of the temporal locations of the labeled events. As shown in Fig. 1, each segment that contained either epileptic or non-epileptic labeled EEG was extracted as a valid segment. According to Fig. 1, segments with one or more epileptic spikes are considered epileptic segments even if they have non-epileptic labels, as well. Thus, one segment had 16 electrodes and 512 samples (1.024 s). Z-score normalization was used with the mean value and standard deviation for each segment.

**Figure 1.**
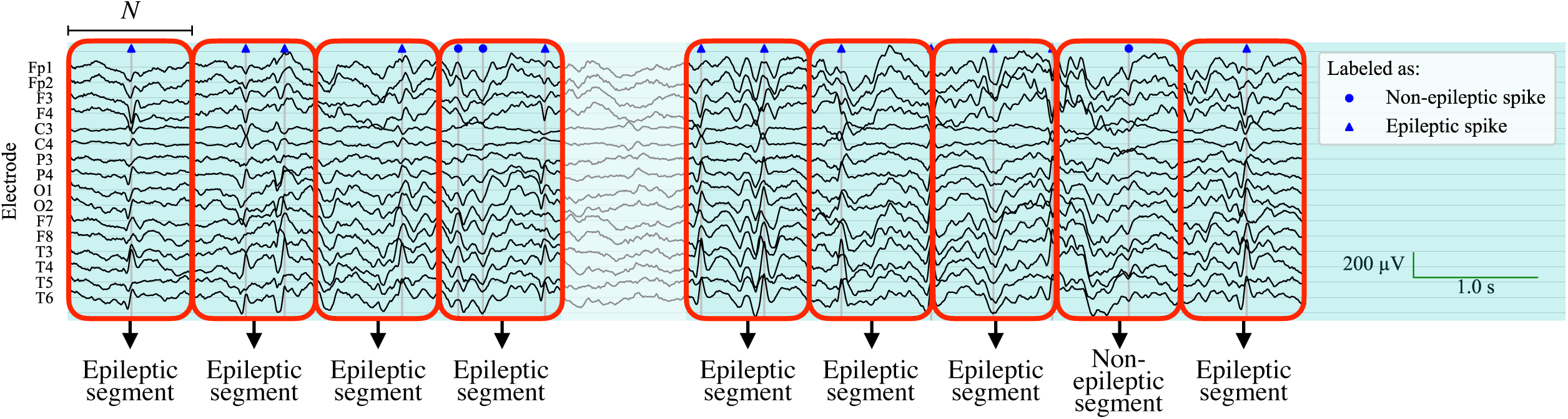
Example of generating segments from the EEG recordings with supervised labels. If a segment contains two distinct labels, it is deemed an epileptic segment. Unlabeled segments are discarded owing to the possibility that unlabeled EEG segments may have epileptic spikes, they are discarded. *N* is the length of the segment and is set to 512 (1.024 s).

### 2.2. Separable Convolution

A layer known as separable convolution, which independently convolves the temporal and spatial directions, should be inserted at the beginning of the network for the analysis of multi-electrode EEG with NNs. This involves frequency filtering for the temporal convolution and electrode coupling for the spatial convolution (i.e., montage optimization). Given that it has a separate kernel for each dimension, this convolutional layer has much smaller parameters than a standard two-dimensional convolutional layer. The first model to use this layer for EEG analysis is EEGNet [21], which has been successful in the field of brain-computer interface. Huang et al. [22] also reported the effectiveness of the EEGNet-based model in their study of EEG-based emotion classification tasks. Using an extended model of EEGNet, Shoji et al. [23] successfully identified abnormal EEG durations indicative of patients with juvenile absence epilepsy. Furthermore, SpikeNet [12] uses a separable convolution to detect epileptic spikes. As these results show, the separable convolution in multi-electrode EEG analysis is effective, particularly in extracting features along the electrode direction.

### 2.3. Self-Attention Mechanism

Accurately detect epileptic spikes, even in randomly extracted EEG segments, without using candidate detectors. However, to train a deep learning model with many parameters [17], a significant amount of training data are needed. We focused on the SA mechanism to efficiently analyze the waveform of interest. The SA mechanism is expected to automatically skip redundant data in time-series signals [18]. Using the dot-product attention [24] is one way to construct SA. That is, the layer based on SA calculates three hidden features, ***Q***^(*i*)^ ∈ ℝ^*τ* ×*d*^, ***K***^(*i*)^ ∈ ℝ^*τ* ×*d*^, and ***V*** ^(*i*)^ ∈ ℝ^*τ* ×*d*^, from the input features ***X***^(*i*)^ ∈ ℝ^*τ* ×*d*^ using the three weight matrices, ***W***_*Q*_ ∈ ℝ^*d*×*d*^, ***W***_*K*_ ∈ ℝ^*d*×*d*^, and ***W***_*V*_ ∈ ℝ^*d*×*d*^, as follows:

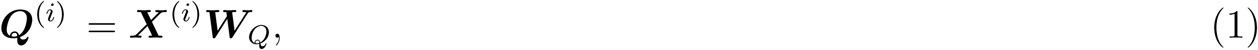

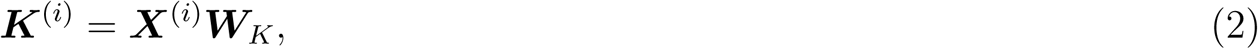

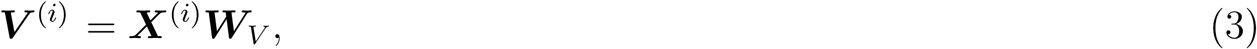

where *i, τ*, and *d* are the index of EEG segments, the temporal length of the input feature, and the number of feature electrodes, respectively. Then, using the softmax function and one weight matrix ***W***_*O*_ ∈ ℝ^*d*×*d*^, the output of the SA layer ***Y***^(*i*)^ ∈ ℝ^*τ* ×*d*^ is obtained as follows:

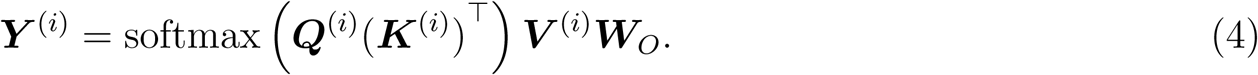

As (4) shows, the softmax function of the matrix product between ***Q***^(*i*)^ and ***K***^(*i*)^ acts to represent the level of interest within the input feature itself. Further, it is expected that multiplying the softmax using the weighted input feature ***V***^(*i*)^ extracts meaningful EEG locations for the prediction. Here, the SA layer can be constructed as a single layer using these calculations.

### 2.4. Satelight: A self-attention-based lightweight model

In this study, we propose a model that can be trained using multi-electrode EEG segments and labels that indicate whether the spikes are epileptic. Fig. 2 illustrates the architecture of the proposed Satelight. As shown in Fig. 2, the model is factorized into the first convolution block and three SA blocks. The first convolution block combines temporal and spatial convolutions, whose effectiveness in lowering the number of learnable parameters has been reported in [21, 12]. A two-dimensional convolution layer with a kernel size of (*f*_*s*_*/*2, 1) implements the temporal convolution. Because the temporal kernel size is half that of the sampling frequency, it theoretically constitutes a filter with a frequency resolution of 2 Hz [21]. Spatial convolution is performed using a two-dimensional depthwise convolution layer. In this layer, 16 input matrices of size (*N, C*) are independently convolved with *C*-sized kernels. Thus, it is expected that the desired electrodes will be selected. These convolutions feed the following SA blocks with the features that have been spatiotemporally filtered. Therefore, every 8 ms, the SA layer is expected to search for relationships between feature vectors. Finally, the fully connected layer with sigmoid activation is adopted as the output layer for binary classification. For training Satelight, cross-entropy, which is the log-likelihood of the Bernoulli distribution, is employed as the loss function:

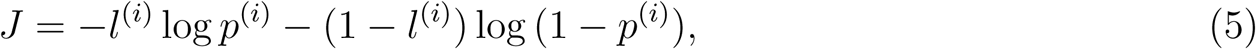

where *l*^(*i*)^ ∈ {0, 1} and *p*^(*i*)^ ∈ [0, 1 denote the annotated label and the model’s prediction of the *i*-th EEG segment, respectively.

**Figure 2.**
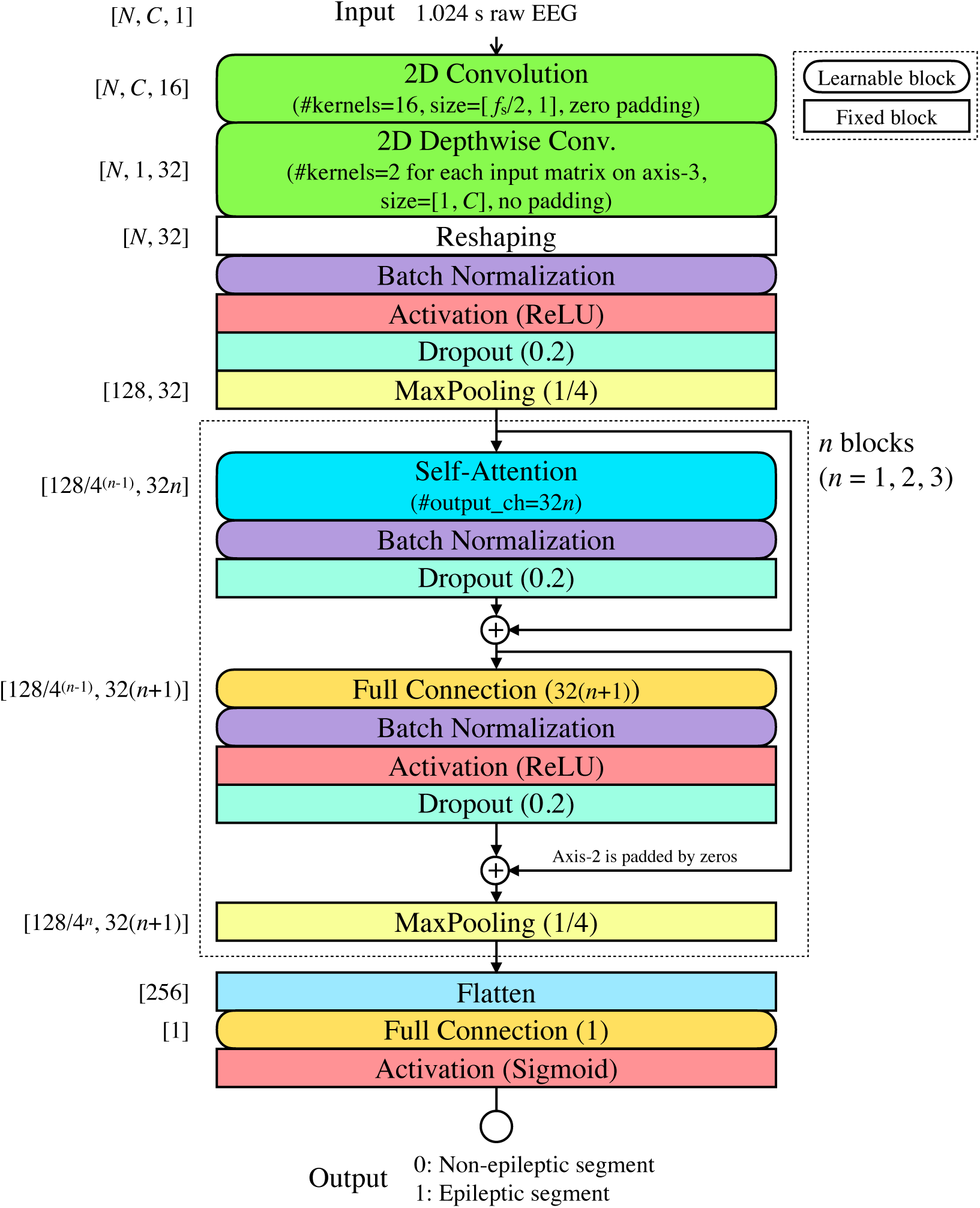
Architecture of the proposed Satelight, where *N, C*, and *f*_*s*_ are the length of the input segment, the number of EEG electrodes, and the sampling frequency, respectively. In this study, *N* = 512, *C* = 16, and *f*_*s*_ = 500.

### 2.5. Experimental implementation

To verify the effectiveness of the proposed Satelight, we conducted a numerical experiment using surface EEGs recorded from patients with epilepsy. As mentioned, this experiment is a binary classification of segments that contain or do not contain epileptic spikes. The following five NN-based models, one statistical property-based model, and one commercially available software are used as comparison models:

- NN-based models:
  − The proposed model (Satelight);
  − Thomas et al.’s model [14];
  − Two models proposed by Jing et al. [12] with the kernel size of all temporal convolutional layers set to:
    ∗ three (SpikeNet3);
    ∗ five (SpikeNet5);
  − Lawhern et al.’s model (EEGNet) [21];
- Statistical property-based model:
  − Janca et al.’s model [25];
- Commercially available software:
  − Persyst13 software [16].

In this experiment, 50 leave-one-patient-out tests were performed using EEG segments from 49 patients as a training set and the remaining segments from one patient as a test set in all possible combinations. The Adam optimizer [26] used the training set to train the models for comparison. Additionally, early stopping [27] was applied using a portion of the training set (namely, the validation set) to suppress overfitting. The validation set consists of 20% randomly selected segments from the training data. Thus, 80% of the segments were used for training in the 49 EEG recordings. The Xavier initializer [28] was used to generate the initial weights for this model. Note that the training process was not conducted on Persyst13 software.

To compare the classification performances, we used four evaluation criteria: the area under the curve (AUC) of the receiver operating characteristic (ROC), F1-value, true positive rate (TPR, sensitivity), and false-positive rate (FPR). F1-value, TPR, and FPR are calculated as follows:

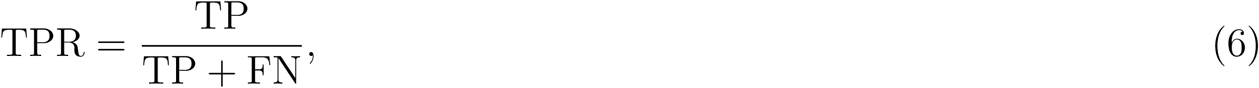

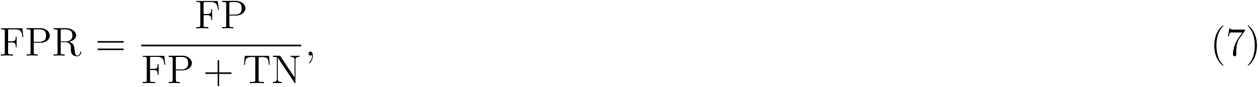

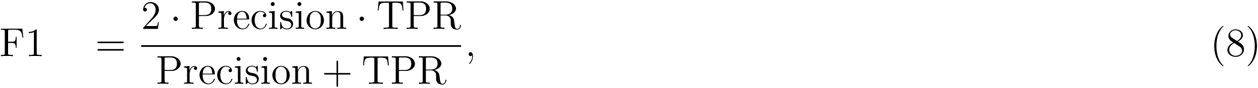

where

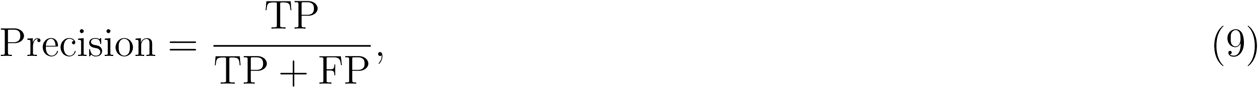

where TP, FP, FN, and TN represent the values of true positive, false positive, false negative, and true negative segments, respectively. The AUC denote the area of the ROC drawn by the TPR and FPR when the classification threshold is changed from zero to one.

This experiment was computed using Python 3.7.6 with Keras [29] and Scikit-learn [30] on a high-performance computer built with an AMD(R) EPYC(TM) 7742 CPU@2.25 GHz, 512 GB RAM, and four NVIDIA(R) A100 GPUs.

## 3. Results

### 3.1. Results of spike detection performance

Table 2 shows the averaged numerical results of the 50 leave-one-patient-out tests. According to Table 2, the proposed method outperformed all evaluation metrics except AUC. Friedman’s one-way analysis of variance (ANOVA) [31] showed that the effects of the models on two metrics—F1 and AUC—were significant (*χ*^2^_F1_(4) = 98.6, *p*_F1_ = 1.95 × 10^*−*20^, *W*_F1_ = 0.499, *χ*^2^_AUC_(4) = 130, *p*_AUC_ = 4.93 × 10^*−*27^, and *W*_AUC_ = 0.648, where *W* denote Kendall’s Coefficient of Concordance [31]). To better understand the effectiveness of the models, we performed a Bonferroni *post-hoc* test [31] because the main effect of the models has been observed. Fig. 3 presents the numerical results and the statistical analysis of the variance of 50 leave-one-patient-out tests. According to Fig. 3, the performance of SpikeNet and Satelight was found to be statistically different from that of EEGNet and Thomas et al.’s model. Although no statistical difference was found between SpikeNet and Satelight, SpikeNet consists of 24 convolutional layers.

**Table 2.**
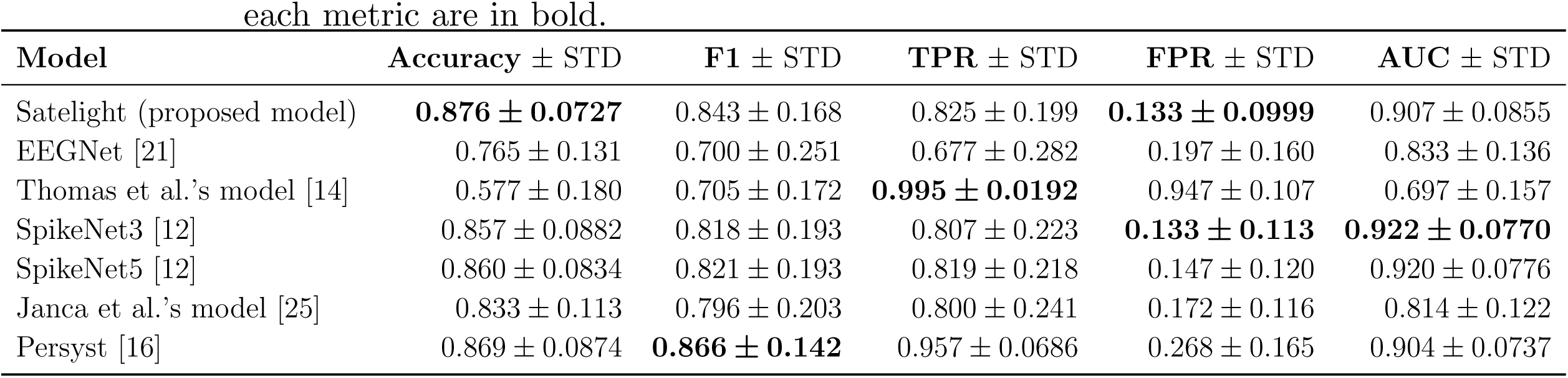
Numerical results of the 50 leave-one-patient-out tests. The best scores for each metric are in bold.

**Figure 3.**
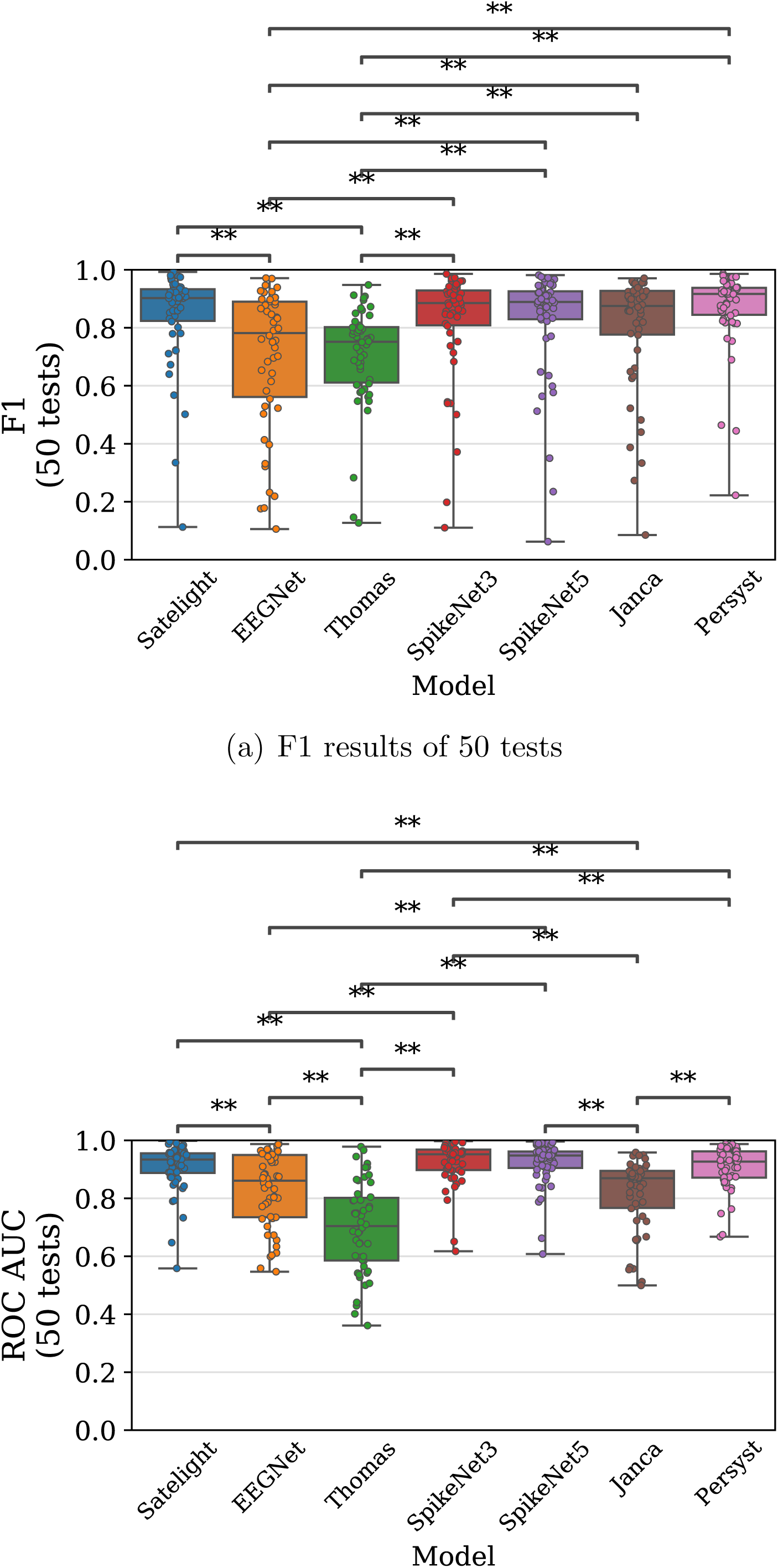
Visualized results in understanding the differences between models. The results of 50 leave-one-patient-out tests with 49 training and one test patient are plotted (excluding Janca et al.’s model and Persyst). For Janca et al.’s model and Persyst, which do not require training, 50 results for each patient are plotted. Statistical significance is indicated by an asterisk (*: *p <* 0.05, **: *p <* 0.01).

### 3.2. Results of self-attention layer behavior

In this section, we discuss an analysis of the behavior of the SA layer in Satelight. Figs. 4 and 5 show heat maps of the first SA layer’s output (the size is [128, 32]), where the SA structure is illustrated in Fig. 2, and the upper EEG segments are input. As shown in Fig. 4, the epileptic spikes appeared between 0.1–0.2 s. The SA output also responded strongly in this range. Similarly, as shown in Fig. 5, the SA responded strongly to the epileptic spikes around 0.7 s. In this case, it did not respond to the artifacts seen around 0.3s.

**Figure 4.**
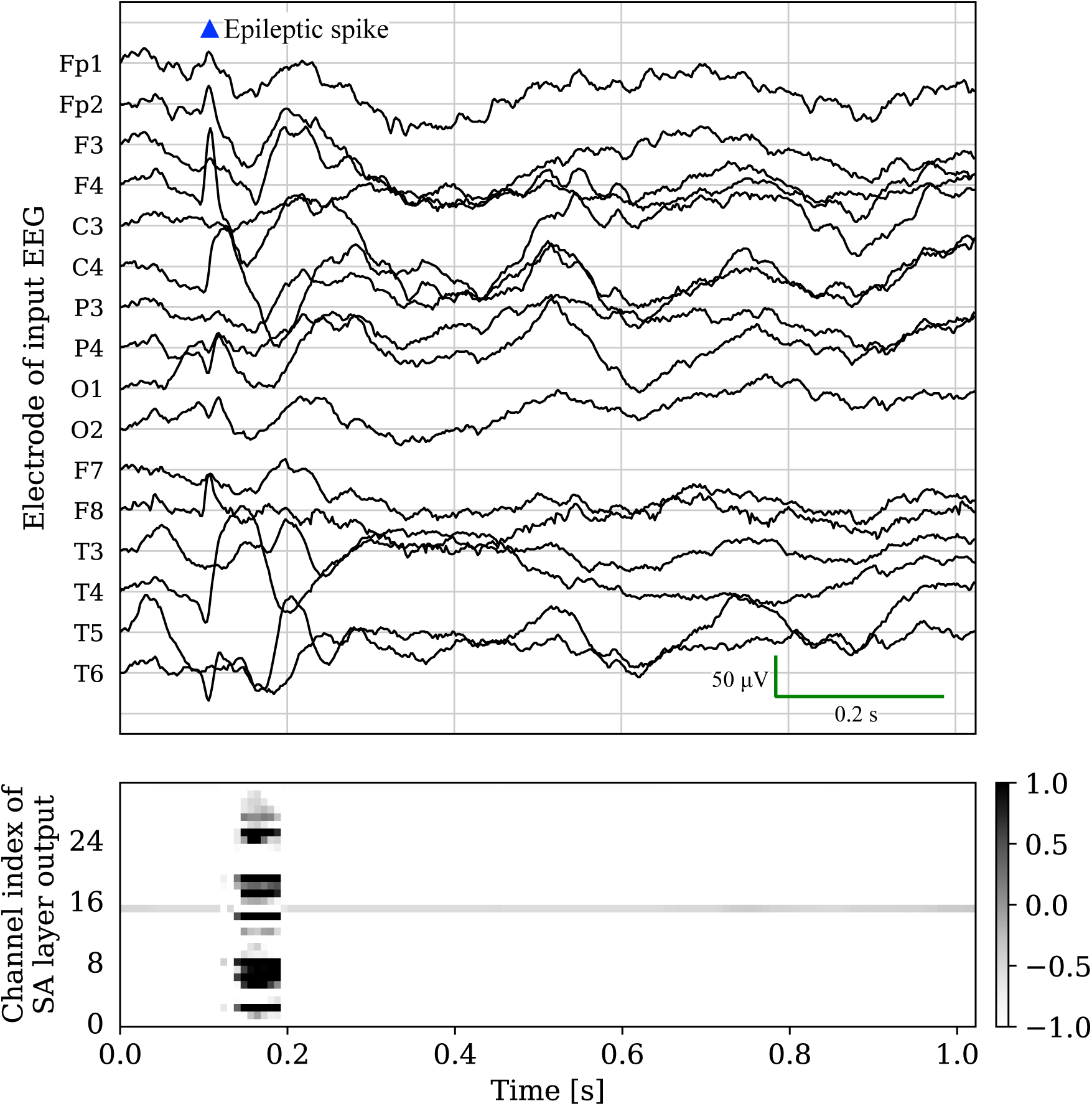
Example of a hidden feature map of the first SA layer (bottom) when predicting an epileptic segment (top). For better visualization, the 128 temporal points in the feature map are stretched as 1.024 s. The peak waveform between 0.1–0.2 s is an epileptic spike.

**Figure 5.**
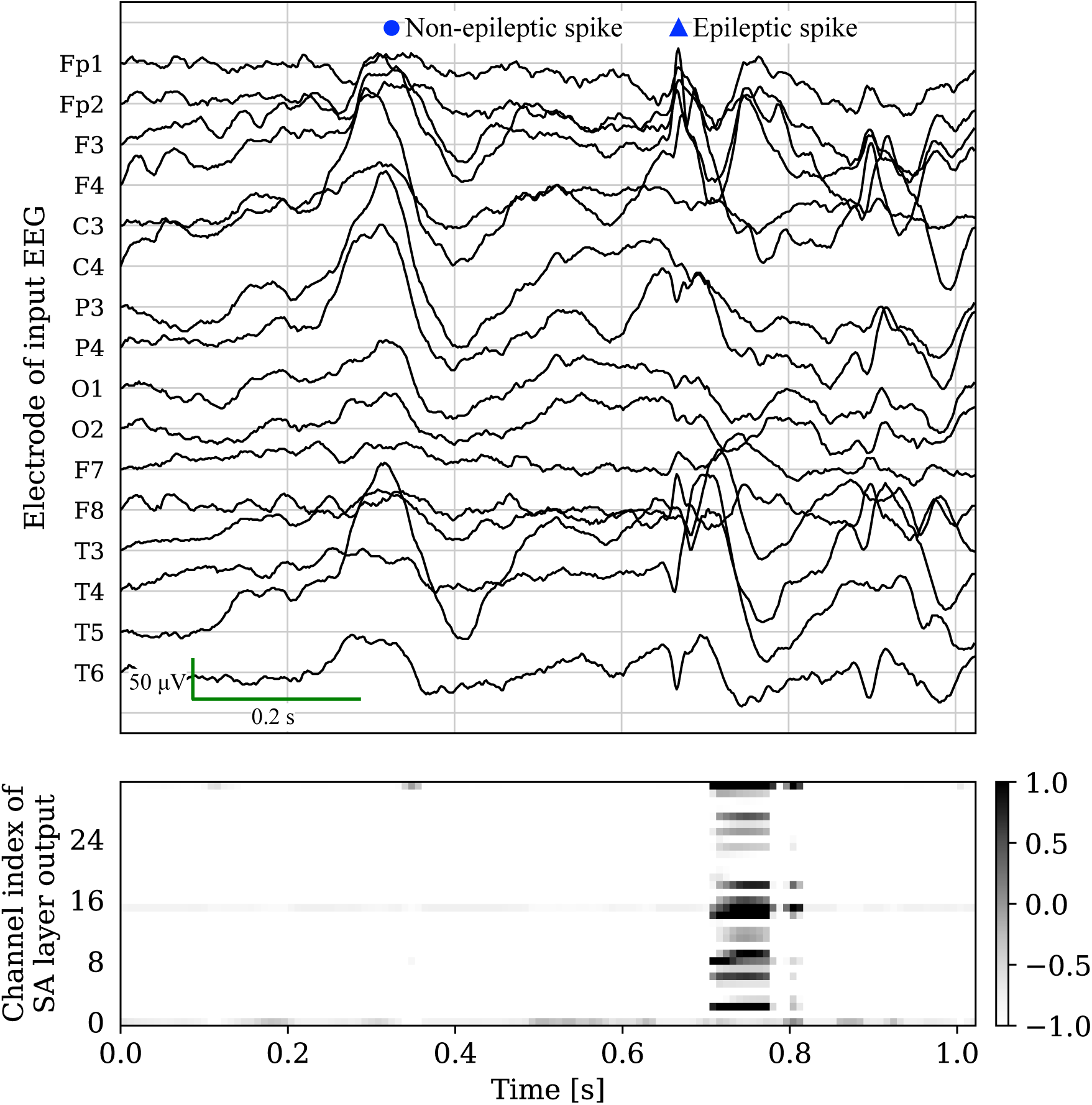
Example of a hidden feature map of the first SA layer when predicting an epileptic segment with extreme artifacts at approximately 0.3 s. For better visualization, the 128 temporal points in the feature map are stretched as 1.024 s. The peak waveform at around 0.7 s is an epileptic spike.

## 4. Discussion

### 4.1. Comparison of the number of parameters in the models

Table 3 and Fig. 6 show the number of parameters used in the numerical experiment for the compared models. EEGNet is a model with few parameters; however, as shown in Table 2, its detection performance is not as high as that of SpikeNet and Satelight. The model by Thomas et al., which had the highest number of parameters, produced a high FPR, as shown in Table 2. This might be because, although it is a deep network, it does not extract features in the spatial direction. Compared with these two models, SpikeNet is more stable in detecting epileptic spikes (accuracy = 0.860 and F1 = 0.821 using SpikeNet5). A combination of deep convolution layers in both the spatial and temporal directions could have been achieved used to achieve this. Moreover, even though Satelight performed better at classification than SpikeNet5 (accuracy = 0.876 and F1 = 0.843), it can be constructed with one-tenth as many parameters. This implies that the deep convolutional layer may be redundant. Therefore, using the SA layer for epileptic spike detection is effective.

**Table 3.** Number of parameters of the comparison models.

| Model | No. of parameters | Ratio with Satelight |
| --- | --- | --- |
| Satelight (proposed model) | 83,905 | $\times 1.00$ |
| EEGNet [21] | 3,061 | $\times 0.0365$ |
| Thomas et al.’s model [14] | 16,387,578 | $\times 195$ |
| SpikeNet3 [12] | 518,658 | $\times 6.18$ |
| SpikeNet5 [12] | 848,386 | $\times 10.1$ |

**Figure 6.**
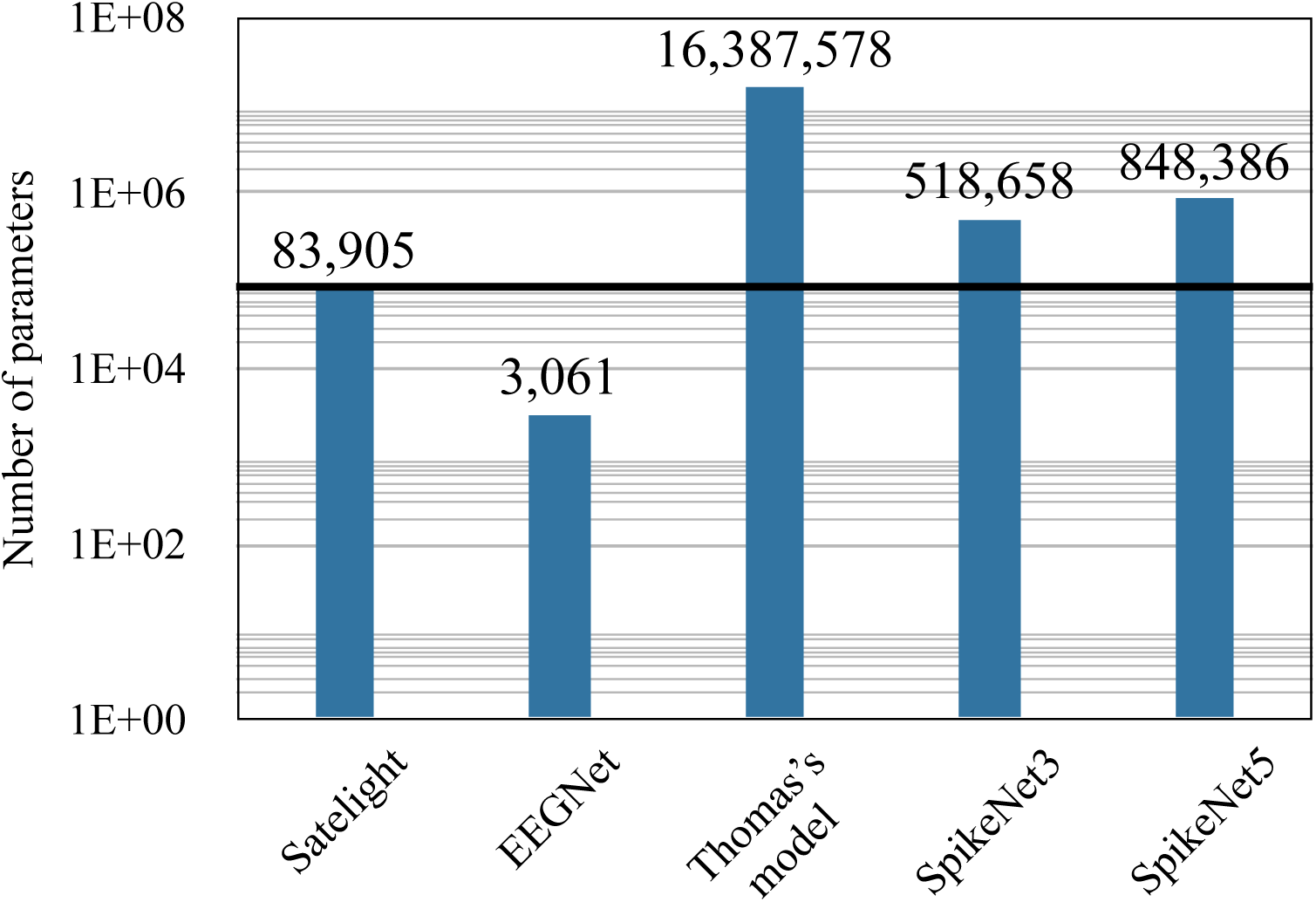
A visual comparison of the model parameters’ numbers. The proposed model (Satelight) is lighter than the other models except for EEGNet, even though it achieves the highest accuracy of all models.

### 4.2. Analysis of self-attention layer behavior

The SA layer behavior analysis results, such as the strong response during epileptic spike and no response on artifacts, as shown in Figs. 4 and 5 may imply that SA operates independently of the EEG amplitude. Moreover, these heat maps demonstrate that the SA responses are slightly delayed (≈ 200 ms) from the onset of the spike. This delay appears to be caused by specialists’ focus on the positive phase reversal following the spike [32]. According to the findings, this experiment, which classifies epileptic spikes and artifacts, does not depend on the EEG near the beginning of the event. Therefore, the behavior of the SA may change if the task involves identifying specific epileptic spike types, such as a spike–wave or sharp–wave.

### 4.3. Dataset collection

Generally, a dataset should include a variety of age, gender, and epilepsy types, as well as a large amount of data and accurate annotations. In this study, as many as 50 EEG recordings were collected for reliable validation. Additionally, the Z-score normalization was applied as preprocessing step to the EEG segments to reduce amplitude differences among individuals. Note that spectralization as preprocessing step is also available [33], however, this paper did not adopt it owing to the increase in configurations to be adjusted—such as window size and window function—and the increases in input dimensions and learnable model parameters. We consider that the proposed model trained on this dataset has high versatility for typical epileptic spike detection. To further improve versatility, the correctness of annotations must be examined. In this study, all samples were annotated at the sole discretion of the annotators without consulting other annotators. That is, the annotation process may contain human errors. The SpikeNet detection result in our experiment (AUC = 0.92) was less accurate than the original result (AUC = 0.98) reported by Jing et al. [12]. One possible reason is that only samples that six or more annotators classified as epileptic were used in their evaluation.

In this study, the age range of the dataset was limited to children. Although it has not been reported that the typical form of epileptic spikes varies with age and gender; however, it is essential to collect patient data from a variety of age groups for reliable validation. Additionally, to build a more practical model, other epileptiforms, such as quasi-periodic spikes and poly-spike, must be annotated. To solve these limitations, it is desirable to develop a framework for more efficient data collection and labeling.

### 4.4. Segment extraction for dataset construction

In many spike detection studies, after detecting candidate spikes and annotating them with labels for the dataset construction, the EEG signals are segmented out based on the labeled spikes. Candidate spikes are typically found using simple signal processing methods and expert selection directly from the EEG. Single-electrode EEG studies use the former method [8, 10, 34]. The latter method is used for both single and multi-electrode EEG studies [12, 13, 14]. Because the candidate spikes are not uniquely defined, there is currently no simple segment method for multi-electrode EEG extraction. For example, detecting peak spikes based on amplitude is challenging becasue spikes do not always occur at precisely the same time over multiple electrodes. Furthermore, depending on epilepsy symptoms, the number of electrodes in which spikes appear varies; thus, the rules for candidate detection are likely to be complex.

However, if segments are extracted based on the expert’s selected spikes, the validation data appear to depend on the expert’s decision even, and then a data set is constructed using these segments. Therefore, for sufficient validation, the expert’s decision must be removed from the factors in the validation phase. Following this justification, the dataset for this study was created using the following methodology:

i. Split the EEG recordings as segments at fixed intervals from the beginning.
ii. Indicate on labels whether the segment contains epileptic spikes.

The method for extracting segments of the validation data using this procedure was superior to the method relying on expert selection. The disadvantage of this method is that the location of spikes varies for each segment; thus, a more effective method for segment extraction or identifying characteristic waveforms for multi-electrode EEG may be necessary. In this study, the SA mechanism, as described in Section 3.2, rather than the segment extraction method, successfully mitigated this limitation.

### 4.5. Implementation for practical use

In this study and numerous others, datasets were built using pairs of EEG segments and corresponding labels, then divided into training and test sets for verification [8, 10, 34, 12, 13, 14]. This validation will be sufficient if spike detection can be considered a simple classification task. However, this validation method using a pre-constructed dataset is a limitation of these studies because the raw EEG recordings are not segmented. Therefore, ideally, all EEG recordings should be fully labeled by the experts without mistakes or omissions. Alternatively, detecting spikes from the entire EEG recordings and then evaluating these predictions posteriorly, rather than constructing the test set, is preferable. In other words, for practical use, a fully automated method for detecting spikes in raw EEG recordings, including EEG segmentation steps, and evaluating them appropriately is urgently required.

## 5. Conclusion

We propose a lightweight epileptic spike detection model that employs the SA mechanism. The number of parameters in this model is small compared with that of the other state-of-the-art deep NN-based models. Nevertheless, the model achieved high detection performance (accuracy = 0.876 and FPR = 0.133). Furthermore, an exploration of the hidden parameters showed that the model automatically paid attention to the characteristic waveform locations of interest in the EEG. This would significantly contribute to the development of an explainable NN.

## Acknowledgements

The authors thank Ms. Meiko Sakurai and Ms. Junko Hirota for their help with annotation as well as Dr. Miao Yao and Mr. Kazi Mahmudul Hassan for an insightful discussion on technical analysis. This work was supported by JST CREST Grant Number JPMJCR1784. We would like to thank Editage (www.editage.com) for English language editing.

